# Evolution of dosage compensation does not depend on genomic background

**DOI:** 10.1101/2020.08.14.251801

**Authors:** Michail Rovatsos, Lukáš Kratochvíl

**Affiliations:** Department of Ecology, Faculty of Science, Charles University, Prague, 12844, Czech Republic

**Keywords:** *Anolis*, dosage compensation, gene expression, sex chromosomes, softshell turtles, transcriptome

## Abstract

Organisms evolved various mechanisms to cope with the differences in the gene copy numbers between sexes caused by degeneration of Y and W sex chromosomes. Complete dosage compensation or at least expression balance between sexes was reported predominantly in XX/XY, but rarely in ZZ/ZW systems. However, this often-reported pattern is based on comparisons of lineages where sex chromosomes evolved from non-homologous genomic regions, potentially differing in sensitivity to differences in gene copy numbers. Here we document that two reptilian lineages (XX/XY iguanas and ZZ/ZW softshell turtles), which independently co-opted the same ancestral genomic region for the function of sex chromosomes, evolved different gene dose regulatory mechanisms. The independent co-option of the same genomic region for the role of sex chromosome as in the iguanas and the softshell turtles offers a great opportunity for testing evolutionary scenarios on the sex chromosome evolution under the explicit control for the genomic background and for gene identity. We showed that the parallel loss of functional genes from the Y chromosome of the green anole and the W chromosome of the Florida softshell turtle led to different dosage compensation mechanisms. Our approach controlling for genetic background thus does not support that the variability in the regulation of the gene dose differences is a consequence of ancestral autosomal gene content.

## Introduction

Sex chromosomes evolve from a pair of autosomes, where one chromosome acquires a sex-determining locus. The regions around this sex-determining locus often stop recombination with their respective homologous regions on X or Z chromosomes (Muller, 1918; Ohno, 1967; reviewed in Charlesworth, Charlesworth, & Marais. 2005), potentially due to sexually antagonistic selection, which leads to the accumulation of alleles beneficial for one sex but detrimental to the other in the vicinity of the sex-determining locus. Over time, the cessation of recombination triggers changes mainly in the Y and W chromosomes, including the accumulation of deleterious mutations and extensive degradation of the gene content. Thus, the differentiation of sex chromosomes leads to unequal numbers of functional copies of many genes between the sexes. These differences have to be in some way handled at the cellular level, as the protein production in a cell is generally affected by the number of transcribed gene copies, and cell physiology and differentiation require proper stoichiometric ratios of interacting proteins (Birchler, Riddle, Auger, & Veitia, 2005; Zhag & Oliver, 2007; Birchler, 2014; Dürrbaum & Storchová 2016). Different lineages evolved distinct mechanisms to cope with the gene copy disequilibrium. Some lineages evolved dosage compensation, the epigenetic mechanism which restores the expression of the X- or Z-specific genes in the heterogametic sex to the ancestral autosomal levels (Muller, 1918; Ohno, 1967; Brockdorff & Turner, 2015).

Despite the common features of the differentiation process of sex chromosomes, it was suggested that there is a dichotomy in the gene dose regulatory mechanisms between male heterogamety (XX/XY) and female heterogamety (ZZ/ZW systems). Complete dosage compensation or at least parity in the expression of the X- or Z-specific genes between sexes (this parity is also referred to as “dosage balance” in the expression levels by some authors, e.g. Gu & Walters, 2017) was often found in lineages with XX/XY sex chromosomes. The term dosage balance refers to the situation where the expression of the Z/X-specific gene is equal between the two sexes, regardless of the ancestral expression level of the same gene when it was autosomal (Gu & Walters, 2017). Such mechanisms have been documented in several insect lineages, nematode worms, therian mammals and the green anole (reviewed in Gu & Walters, 2017). However, lack of dosage balance in the expression of X-specific genes was found in three lineages with male heterogamety: the three-spined stickleback, the platypus and the brown basilisk (Gu & Walters, 2017; Acosta et al., 2019; Nielsen et al., 2019). A lack of dosage balance seems to be common in lineages with female heterogamety, where it was documented in parasitic bloodflukes, tonguefish, caenophidian snakes, the Komodo dragon and birds (Mank, 2009; Vicoso, Emerson, Zektser, Mahajan, & Bachtrog, 2013; Gu & Walters 2017; Picard et al., 2018; Rovatsos, Rehák, Velenský, & Kratochvíl, 2019). The single exception is lepidopteran insects, where dosage balance was found, but the level of expression of Z-specific genes is likely lower than the ancestral state (Huylmans, Macon, & Vicoso, 2017). However, from the above list of taxa it is evident that our knowledge of gene dose regulatory mechanisms is limited to comparisons of a small number of lineages with highly dissimilar general biology and complexity of body plans and genomes. Moreover, the comparison between differentiated sex chromosomes under male and female heterogamety can be strongly confounded by the non-homology of sex-linked genes among these lineages and consequently, the different tolerance to copy variation of dosage sensitive genes, located in sex chromosomes. By a dosage sensitive gene, we refer to any gene where a change in gene dosage (e.g. copy number variation) causes a phenotypic effect, no matter the precise mechanism of dosage sensitivity (reviewed e.g. in Rice & McLysaght, 2017; Pessia, Engelstädter, & Marais, 2014; Zimmer, Harrison, Dessimoz, & Mank, 2016).

Our study suggests a solution to these problems. We compared the gene dose regulatory mechanism in two reptile lineages (i) with opposite heterogamety and (ii) ancient highly differentiated sex chromosomes, which (iii) independently evolved from the same genomic region: the iguanian green anole (*Anolis carolinensis*) with male heterogamety (Alföldi et al., 2011) and the Florida softshell turtle (*Apalone ferox*) with female heterogamety (Rovatsos, Praschag, Fritz, & Kratochvíl, 2017). Both lineages co-opted the same genomic region for the function of sex chromosomes containing genes with orthologs linked to chicken (GGA) chromosome 15 (GGA15) (Alföldi et al., 2011; Rovatsos et al., 2017; Marin et al., 2017). Twelve families of iguanas including anoles share the same X-specific gene content, which documents that differentiated XX/XY chromosomes homologous to those of the green anole were present already in the common ancestor of iguanas living at least *c*. 70–95 million years ago (Rovatsos, Pokorná, Altmanová, & Kratochvíl, 2014a; Altmanová, et al., 2018). In the softshell turtles, the differentiated ZZ/ZW sex chromosomes are stable and can be traced back to the last common ancestor of the extant species, as 10 trionychid species covering the phylogenetic diversity of the family share the same Z-specific genes (Rovatsos et al., 2017). This evidence suggests that trionychid sex chromosomes are likely older than 120 million years (Rovatsos et al., 2017). The long-term stability of sex chromosomes in both lineages should have assured sufficient time for the emergence of an optimal gene dose regulatory mechanism.

The gene content of the X chromosome in the green anole has been extensively identified (Alföldi et al., 2011; Rovatsos, Altmanová, Johnson Pokorná, & Kratochvíl, 2014b; Marin et al., 2017), the Y chromosome is highly degenerated and the complete dosage compensation was recently reported (Marin et al., 2017; Rupp et al., 2017). The dosage compensation in the green anole is reached by up-regulation of genes linked to X chromosome in males. This careful regulation suggests that the genes linked to the X chromosome should be highly dosage sensitive. We therefore predicted that we would find a similar mechanism in the turtle, where the Z chromosome was derived from the same ancestral autosome and the W is also highly degenerated (Rovatsos et al., 2017). Here, we test this hypothesis by determining the Z-specific genes and the sexual differences in their expression in the Florida softshell turtle *A. ferox* and by comparing the expression pattern of the same orthologous genes which are X-specific in the anole and at the same time Z-specific in the turtle.

## Material and methods

### Studied material

Two males and two female**s** of *A. ferox* were obtained from a pet shop in order to collect blood samples for genetic and genomic analyses. Genomic DNA was extracted from all samples using the DNeasy Blood and Tissue Kit (Qiagen, Germany). Total RNA was extracted using TRIzol reagent (Invitrogen, Carlsbad, CA, USA) according to the manufacturer’s protocol.

### Illumina sequencing (DNA-seq, mRNA-seq) and bioinformatic analyses

Genomic DNA from one male and one female of *A. ferox* were sequenced at high coverage (approx. 50x) by Novogene (Cambridge, UK) in Illumina HiSeq2500 platform, with 150 base pairs (bp) pair-end option (DNA-seq). Libraries from total RNA of two males and two females of *A. ferox* were constructed by GeneCore (EMBL, Heidelberg, Germany) (mRNA-seq). The barcoded stranded mRNA-sequencing libraries were prepared using the Illumina TruSeq mRNA v2 sample preparation kit (Illumina, San Diego, CA, USA) with poly-A mRNA enrichment, implemented in the liquid handling robot Beckman FXP2. 84 bp fragments were sequenced unidirectionally in the pooled libraries using the Illumina NextSeq 500 platform. The raw Illumina reads from both DNA-seq and mRNA-seq of all individuals are deposited in Genbank (BioProject PRJNA608206, accession numbers SRR11149095-SRR11149100).

Adapters and low-quality bases from raw reads were trimmed by Trimmomatic (Bolger, Lohse, & Usadel, 2014) and Geneious v. R7.1 (Kearse et al., 2012) using “trim” utility with default parameters. Reads shorter than 50 bp were removed, resulting in the final dataset of 658-731 million reads per specimen for the DNA-seq and 35-78 million reads per specimen for the mRNA-seq. Trimmed reads were checked in FASTQC (Andrews 2010) and MULTIQC (Ewels, Magnusson, Lundin, & Käller, 2016).

In ZZ/ZW sex determination systems with a highly degenerated W chromosome, Z-specific genes have half copy numbers in the genomes of ZW females in comparison to ZZ males. These differences in the copy numbers of Z-specific genes between sexes are detected by the differences in coverage of the reads from DNA sequencing in Illumina HiSeq platform (e.g. Vicoso et al., 2013; Picard et al., 2018). Z-specific loci are expected to have half read coverage in ZW females in comparison to ZZ males, while autosomal and pseudoautosomal loci should have equal read coverage in both sexes. We used this approach for identification of Z-specific genes in *A. ferox*. Trimmed DNA-seq reads from a male and a female were independently mapped to a reference dataset of 174,456 exonic sequences previously published in the genome project of the Chinese softshell turtle, *Pelodiscus sinensis*, the closest related species to *A. ferox* with a well-annotated genome (Wang et al., 2013) using Geneious v. R7.1 (for parameters see Table S1). The read coverage of each exon was extracted and the average coverage for an individual gene was calculated in each specimen. We normalized the coverage of each gene for the total number of assembled reads per specimen (see Vicoso et al., 2013). Subsequently, we calculated the ratio of female to male read coverage for each gene.

Trimmed mRNA-seq reads from a single female were assembled *de novo* with Trinity (Grabherr et al., 2011), resulting to 165,925 putative transcripts. The assembled transcripts were compared to the reference transcriptome of *Pelodiscus sinensis* (Wang et al., 2013) using BLAST (Altschul, Gish, Miller, Myers, & Lipman, 1990). 51,045 transcripts of *A. ferox* with higher than 70% similarity spanning over 150 bp of homologous sequences in *Pelodiscus sinensis* were used as the reference transcriptome for further analyses. The Illumina reads from all individuals were mapped independently to this reference transcriptome using Geneious v. R7.1 (for parameters see Table S1). We filtered out all loci not expressed in at least one individual or with transcript length less than 500 bp. To avoid pseudoreplications at the gene level, the subsequent analyses were done using just the longest transcript per gene. We assigned genes to putative syntenic blocks according to chromosome position of their orthologous genes in the chicken genome (http://www.ensembl.org). This procedure is substantiated by the high level of conservation in gene synteny between chicken and turtles (O’Connor et al., 2018). Furthermore, the chicken has one of the best assembled genomes among sauropsids at the chromosome level, facilitating cross-species comparisons. We used this procedure to test whether the region containing Z-specific genes in *A. ferox* is indeed syntenic to GGA15 and thus to the X chromosome of the green anole as previously stated (Rovatsos et al., 2017).

### Validation of Z-specific gene identification by qPCR

We used qPCR for estimation of the difference in gene copy number between male and female genomes in *A. ferox* to validate Z-specificity in selected genes and thus to further support the accurate identification of the *A. ferox* Z-specific gene content. The detailed methodology of this approach is described in Rovatsos et al. (2014a,b; 2016; 2017; 2019). Primers specific for three Z-linked (*anapc7*, *ccdc92, tmem132d*) and three autosomal control (*adarb2*, *mos, rag1*) genes were previously published for the trionychid turtles by Rovatsos et al. (2017). For the validation we used DNA isolated from three males and three females of *A. ferox*.

### *Test of dosage balance in the expression of Z-specific genes in* A. ferox *and direct comparison to* A. carolinensis

The RPKM expression values were independently calculated for each transcript with average read coverage higher than 10 in each specimen, resulting in a final dataset with expression data from 5,616 genes (Table S2). Subsequently, we computed the average sex-specific RPKMs for each transcript as the mean value from the two females and two males, respectively. We tested for dosage balance in the expression of the Z-specific genes by comparing the female to male ratios in RPKM between Z-specific genes and other genes by Mann-Whitney U test. We log_2_-transformed the ratios to improve the symmetry of the distribution of ratios. The non-parametric test was used as Kolmogorov-Smirnov test showed that the data significantly deviate from normality (p < 0.01). Genes with female to male ratio above 2.0 (in total less than 0.8% of genes) were excluded from the analyses as they likely represent highly female-biased genes; however, their inclusion does not change any interpretation.

Our next aim was to compare the sexual differences in the expression of the Z-specific genes of the turtle and of their X-specific orthologs in the green anole directly on a gene-by-gene basis. We identified X-specific orthologs of the *A. ferox* Z-specific genes with expression data in the green anole in the data from Rupp et al. (2017). We compared the female to male ratios in RPKM of the same genes between the anole and the turtle by Wilcoxon signed-ranks test.

Single copy genes linked to the Z-specific region are hemizygous and their transcripts thus should not have any SNPs in females. We utilize these characteristics in combination with information on read coverage in the male and female genomes to identify Z-specific genes even more reliably. For the conservative test of dosage balance in the expression in the turtle we identified Z-specific genes as the genes without SNPs and with the female to male ratio in read coverage depth lower than 0.7.

## Results

The comparative read coverage analysis was performed in 19,151 genes of *A. ferox*, revealing 568 genes with female to male ratio for read coverage less than 0.7, corresponding to Z-specificity (Table S2, Fig. 1). Among these potential Z-specific genes, we identified 245 genes with known chromosomal position of orthologs in chicken genome. Notably, 220 out of 245 potential Z-specific genes of *A. ferox* have orthologs linked to GGA15, while the remaining 25 genes have orthologs scattered to 16 chicken chromosomes (Table S2). We validated sexual differences in gene copy numbers in two identified Z-specific genes by qPCR, applied to male and female genomic DNA as a template. qPCR revealed the expected pattern of approximately half the number of copies in the female genome in comparison to the male genome in all tested Z-specific genes and equal gene copy number in the control autosomal genes (Fig. S1). These results corroborate that the syntenic block homologous to GGA15 is a part of the Z chromosome in *A. ferox* as previously documented by physical gene mapping in the Chinese softshell turtle, *Pelodiscus sinensis* (Kawagoshi, Uno, Matsubara, Matsuda, & Nishida, 2009) and validated in 10 species of softshell turtles by the comparison of gene copy numbers between male and female genomes (Rovatsos et al., 2017). The analysis of the female to male ratios in DNA-seq read coverage uncovered that not all genes with orthologs linked to GGA15 are necessarily in the Z-specific region of *A. ferox*. In total 32 genes with orthologs linked to GGA15 show female to male ratio in read coverage higher than 0.7, corresponding to their autosomal or pseudoautosomal position, or to poorly differentiated Z- and W-specific alleles in the non-recombining region of the turtle Z and W chromosomes (Table S2).

**Fig 1:**
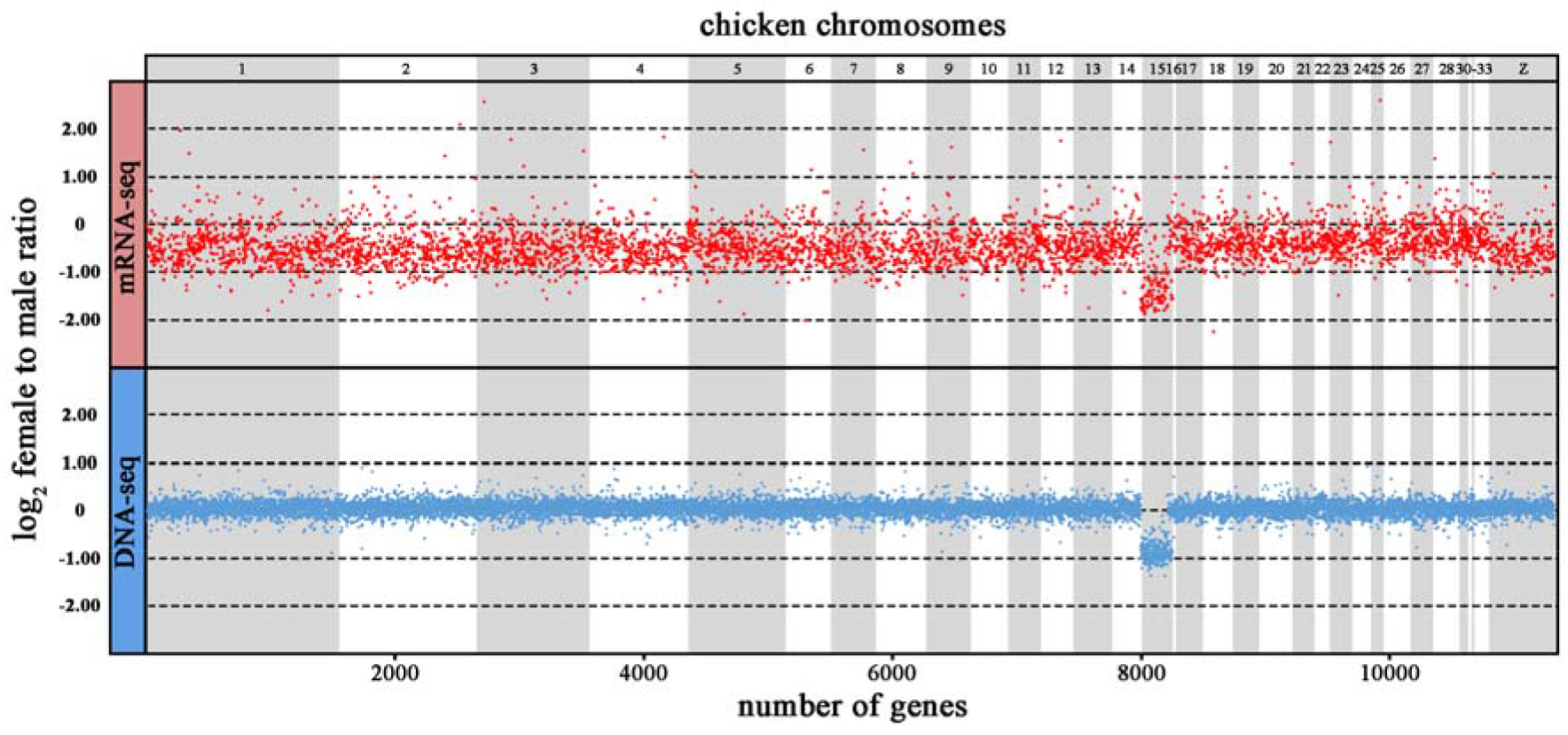
Log_2_-transformed female to male ratios in DNA-seq read coverage (blue) and in expression (RPKM, red) across identified genes of *Apalone ferox*. Each dot corresponds to the f/m ratio from a gene. In the absence of a chromosome-level genome assembly in trionychid turtles, the genes are illustrated according to the position of their orthologs in the chicken genome. Note that the region homologous to chicken chromosome 15 possess much lower ratios in both read coverage depth and RPKM, demonstrating that this part of genome is Z-specific and lacks dosage balance in expression between sexes in most genes in the turtle.

To study whether gene expression is equal in both sexes in the turtle, we analysed our candidate Z-linked genes that had both female to male ratio in read coverage < 0.7. This yielded a total of 118 candidate Z-specific genes in our mRNA-seq dataset. Notably, 102 of them have orthologs on GGA15, which represents 93% of the candidate Z-linked genes with known chromosomal position of orthologs in chicken genome. The female to male ratios in RPKM differ highly between these candidate Z-specific genes and autosomal and psedoautosomal genes (Mann-Whitney U test: U = 55,845, p < 0.0001, n = 5,575), with the median female to male ratio in the expression level being about half of the median of the other genes in our mRNA-seq dataset (Fig. 2). We conclude that there is no dosage balance in the softshell turtle in the Z-specific genes. Expression data were available for 45 orthologues of these genes that show X-specificity in *A. carolinensis* (Rovatsos et al., 2014a,b; Marin et al., 2017; Rupp et al., 2017) (Table S3). Wilcoxon signed-ranks test revealed that these genes have significantly higher female to male ratios in the anole in comparison to the Florida softshell turtle (Z = 6.21, p < 0.0001, n = 51). They are expressed at similar levels in both sexes in the green anole (Fig. 3). The results stayed the same even when a more conservative criterion, i.e. to consider as Z-specific only the genes without SNPs in the turtle, was applied.

**Fig 2:**
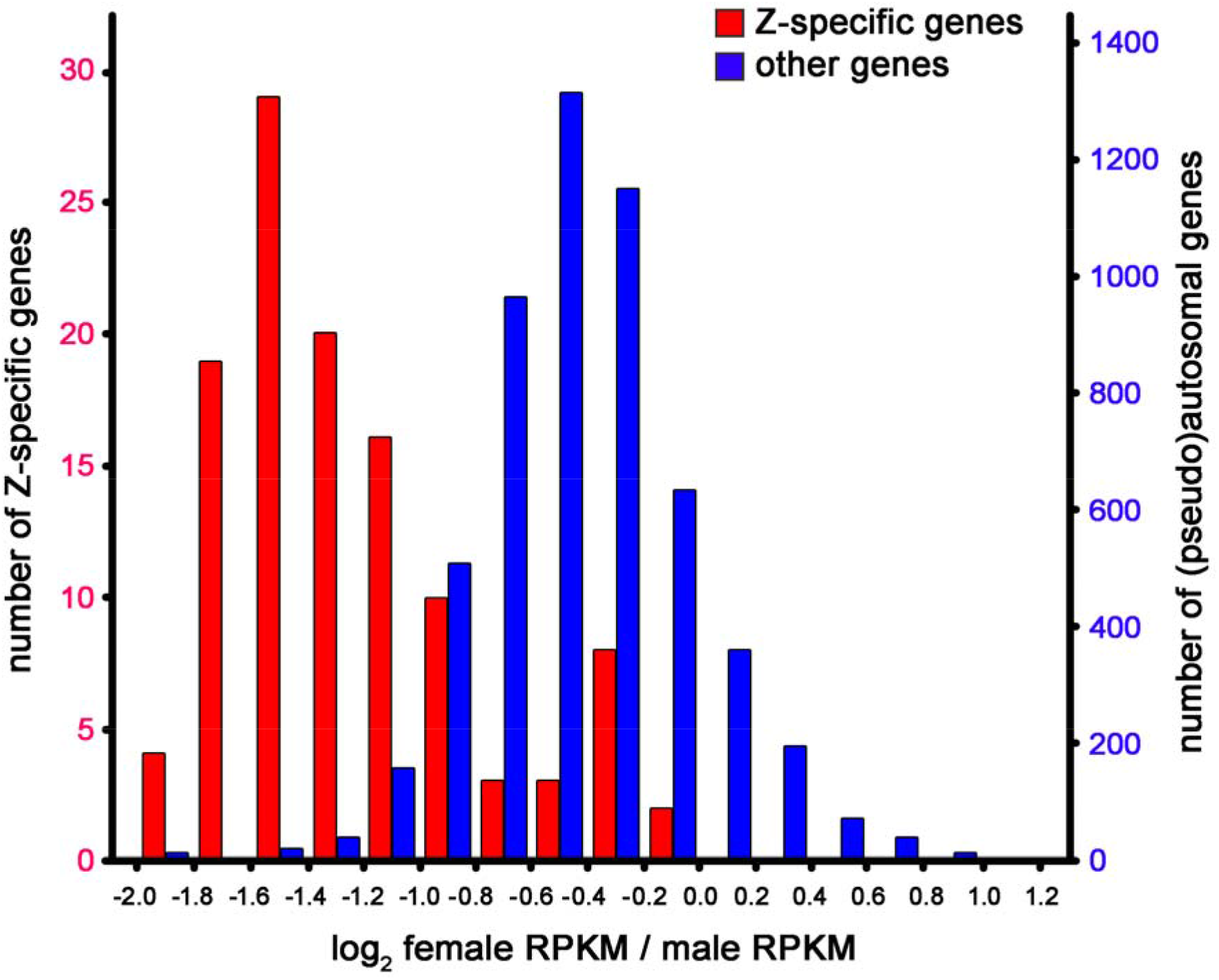
Histogram of the log_2_-transformed female to male ratios in the expression measure (RPKM) for Z-specific genes (red) and other genes (blue) in *Apalone ferox*.

**Fig 3:**
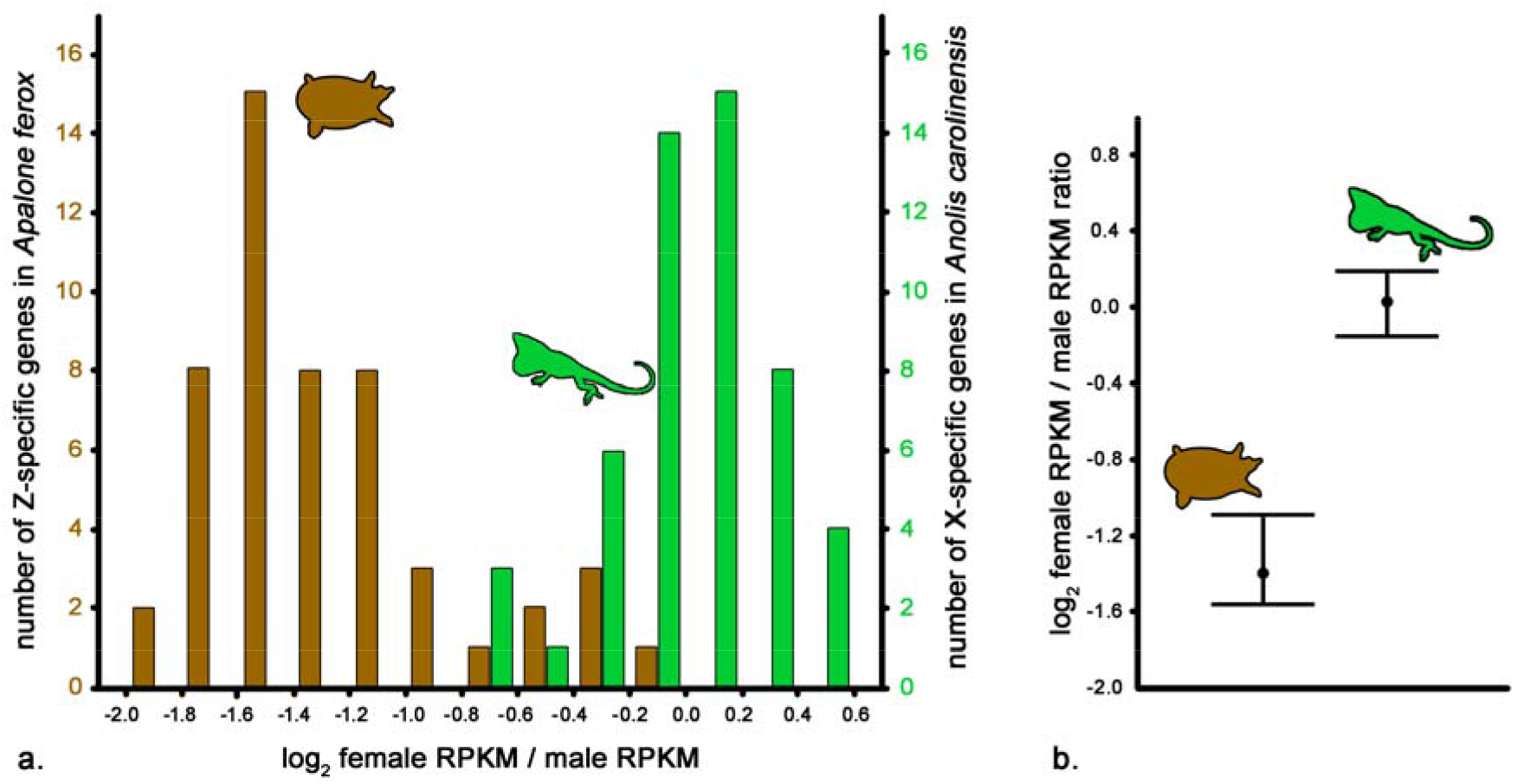
Comparison of the sexual differences in expression of the orthologous genes which are Z-specific in the Florida softshell turtle and X-specific in the green anole. The expression shows mostly dosage balance in the expression between sexes in the lizard but not in the turtle. Histograms (a) and medians and 25^th^ and 75^th^ quartiles (b) of female to male ratios in RPKM are given.

## Discussion

Contrary to our expectations, the sex-specific transcription of the orthologous genes which are X-specific in the green anole and at the same time Z-specific in the softshell turtle differ between the species. The X-specific genes are dosage compensated in the anole, but the expression of the same genes when Z-specific is not equalized between sexes in the turtle (Figs. 2,3). We can thus conclude that the loss of the same functional genes from the Y chromosome of the green anole and the W chromosome of the Florida softshell turtle led to different dosage compensation mechanisms. Our approach controlling for genomic background and gene identity thus shows that the regulation of the gene dose differences is not a consequence of the ancestral gene content of the genomic region now playing the role of sex chromosomes. Moreover, the comparison of the sex-specific expression of the orthologous genes between the turtle and the anole suggests that the dosage compensation of the X-specific genes in the anole does not reflect their sensitivity to gene copy number variation. Orthologs of the dosage-sensitive genes should hence be compensated in the turtle as well, or they should stay in the poorly differentiated regions of the sex chromosomes or be translocated to autosomes. Alternatively, genes linked to sex chromosomes in the anole and the turtle could theoretically change sensitivity to copy number variation during evolution, or sensitive to copy number variation of a gene can be context-dependent (see Deutschbacher et al. 2005; Morrill and Amon 2019). Nevertheless, considering that gene function and expression are generally conserved across vertebrates (e.g. Chan et al. 2009), an hypothetical scenario of mass swift of dose sensitivity seems less likely to explain the differences in gene dose regulation between the Z-specific genes of the green anole and the Z-specific genes of the Florida softshell turtle.

The difference between the anole and the Florida softshell turtle in the dosage compensation mechanisms is in agreement with the often-reported differences between male and female heterogamety. The reasons why these two systems should differ in the dosage compensation mechanisms are not clear and several processes potentially responsible for this dichotomy were suggested (Vicoso and Bachtrog 2009; Mank et al. 2010; Wilson Sayres and Makova 2011; Naurin et al. 2012; Mank 2013; Mullon et al. 2015). Recently, several exceptions from this pattern were reported and after these additions, lineages with male heterogamety are not significantly more likely to possess dosage balance between sexes in the expression of genes linked to sex chromosomes than lineages with female heterogamety (reviewed in Rovatsos et al. 2020).

We hypothesized that the evolution of dosage compensation mechanism might reflect to some extent differences in the function of sex-determining genes. These genes principally work in two ways: sex determination might be controlled either by the copy number of X or Z-linked loci per cell (i.e. gene dosage), or by the presence of a dominant W or Y locus in the genome (Clinton, 1998). The dosage-dependent sex determination can work only in the absence of a mechanism equalizing the expression of the sex-determining locus between sexes, at least in the time when its expression is crucial for sex determination. In contrast, a chromosome-wide regulatory mechanism of the expression of X- and Z-linked genes leading to dosage balance such as heterochromatinization of a single X copy per cell in female mammals (Brockdorff & Turner, 2015), is compatible with the sex determination based on a dominant factor on Y and W chromosomes (e.g. *sry* gene in viviparous mammals) as well. In support, both studied lineages with female heterogamety likely relying on the dosage-dependent mechanism, i.e. birds and caenophidian snakes (Smith et al., 2009; Rovatsos et al., 2018), do not have dosage balance in the expression of Z-specific genes (Ellegren, 2002; Vicoso et al., 2013).

At first sight, two model organisms, the fruit fly *Drosophila melanogaster* and the nematode worm *Caenorhabditis elegans,* represent a contradictory case, since their sex determination primarily relies on the number of copies of the X chromosome, but at the same time they have global dosage compensation achieved by upregulation of the expression of a single X in males. However, dosage compensation in fruit flies and worms is triggered only later in development, and thus does not interfere with the earlier sex-determination mechanisms based on copy numbers (Baker and Belote 1983; Deng et al. 2011; Zanetti and Puoti 2013). These cases illustrate that detailed knowledge on molecular machinery and timing of particular steps will often be needed for testing mechanistic hypothesis on the evolution of gene dose regulatory mechanisms. Currently, our knowledge on the identity and function of sex determining loci is sporadic and restricted mainly to model organisms (Bachtrog et al., 2014; Pan et al., 2017), but we expect that our hypothesis can be tested in future when more evidence will be accumulated. Based on our hypothesis, the presence of dosage-sensitive mechanism of sex determination is more likely in the softshell turtle.

To sum up, we introduce that independent co-option of the same genomic region for the role of sex chromosome, as seen in the iguanas and the softshell turtles, offers a great opportunity for testing evolutionary scenarios on the sex chromosome evolution under the explicit control for the genomic background. Among amniotes, more lineages than the iguanas and the softshell turtles co-opted the same syntenic block for sex chromosomes, as shown for instance by our ongoing research on lacertid lizards and geckos (ZZ/ZW) and therian mammals (XX/XY) (Rovatsos et al. 2016a; 2016b). Future studies should further utilize these excellent systems to explore the convergent/divergent evolution of sex chromosomes.

## Data access

The raw Illumina reads from DNA-seq and mRNA-seq of all studied individuals are deposited into the NCBI BioProject database with ID PRJNA608206 (accession numbers SRR11149095-SRR11149100).

## Acknowledgements

Four anonymous reviewers and the editors made excellent comments to the former version of the manuscript. We are grateful to Vladimír Beneš (Genecore, Heidelberg) for valuable consulting on NGS sequencing, experimental design and data analysis. Computational resources were provided by the CESNET LM2015042 and the CERIT Scientific Cloud LM2015085 under the project “Projects of Large Research, Development, and Innovations Infrastructures”. This study was supported by Czech Science Foundation (project No. 17-22604S). Internal support was provided by Charles University research project PRIMUS/SCI/46 and Charles University Research Centre program (204069).

---

**Fig S1:**
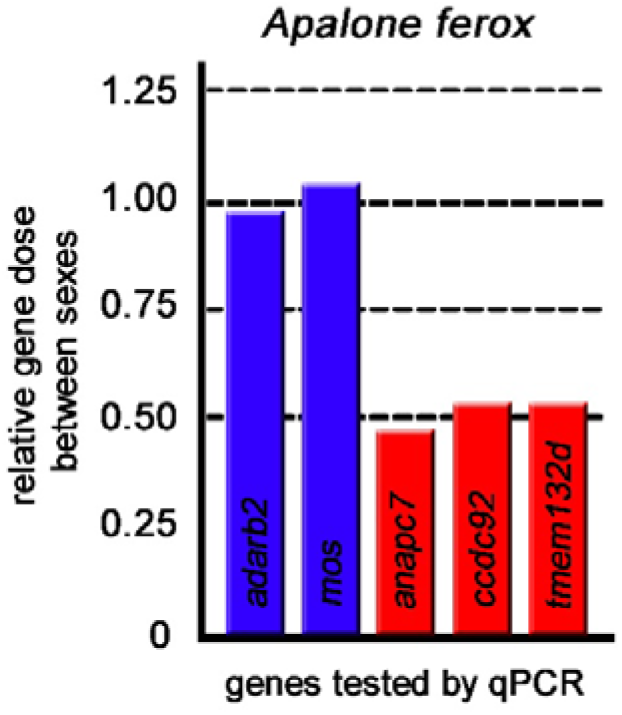
Relative gene dose ratios (r) between females and males for each primer pair for autosomal control (blue) and Z-specific genes (red) in three pairs of *Apalone ferox*. The gene *rag1* was used for normalization of the qPCR values.

## Supplementary information

**Table S1:** Parameters for mapping Illumina mRNA-seq reads in reference transcripts in Geneious v. R7.1.

**Table S2:** List of examined genes from the genome of *Apalone ferox* and the position of their homologous genes to chicken (*Gallus gallus*). Female to male (f/m) ratios are presented for both DNA-seq read coverage analysis and RPKM expression values in *A. ferox*.

**Table S3:** List of 45 orthologous genes which are X-specific in *A. carolinensis* and Z-specific in *A. ferox*. Data for *A. carolinensis* were collected from Marin et al. (2017) and Rupp et al. (2017).

